# Neuromorphic Cytometry: Implementation on cell counting and size estimation

**DOI:** 10.1101/2023.07.06.548044

**Authors:** Ziyao Zhang, Zhangyu Xu, Helen M. McGuire, Chip Essam, Andrew Nicholson, Tara J. Hamilton, Jiayin Li, Jason K. Eshraghian, Ken-Tye Yong, Daniele Vigolo, Omid Kavehei

## Abstract

Flow cytometry is a widespread and high-throughput technology that can measure the features of cells and can be combined with fluorescence analysis for additional phenotypical characterisations but only provide low-dimensional output and spatial resolution. Imaging flow cytometry is another technology that offers rich spatial information, allowing more profound insight into single-cell analysis. However, offering such high-resolution, full-frame feedback can compromise speed and has become a significant trade-off challenge to tackle during development. In addition, the current dynamic range offered by conventional photosensors can only capture limited fluorescence signals, exacerbating the difficulties in elevating performance speed. Neuromorphic photo-sensing architecture focuses on the events of interest via individual-firing pixels to reduce data redundancy and provide low latency in data processing. With the inherent high dynamic range, this architecture has the potential to drastically elevate the performance in throughput by incorporating motion-activated spatial resolution. Herein, we presented an early demonstration of neuromorphic cytometry with the implementation of object counting and size estimation to measure 8 *μ*m and 15 *μ*m polystyrene-based microparticles and human monocytic cell line (THP-1). In this work, our platform has achieved highly consistent outputs with a widely adopted flow cytometer (CytoFLEX) in detecting the total number and size of the microparticles. Although the current platform cannot deliver multiparametric measurements on cells, future endeavours will include further functionalities and increase the measurement parameters (granularity, cell condition, fluorescence analysis) to enrich cell interpretation.

## I. INTRODUCTION

Conventional flow cytometry is a vastly adopted high-throughput technology that is capable of measuring multi-parametric features of cells in a population, including cell count, relative size, granularity, and can be combined with fluorescence detection for additional phenotypical characterisations^3,9^. It can analyse over 10,000 events per second to generate instant feedback on samples^15^. However, such a high-speed approach is limited to lower-dimensional feedback and lacks subcellular resolution^16^. Thus, numerous applications have been developed to optimise various emerging research needs. Imaging flow cytometry (IFC) is one of the remarkable strategies that combines features of high efficiency in flow cytometry with detailed spatial information and fluorescence intensity^2,15^. It can comprehensively visualise cell area with more intricate metrics, including cell morphology, texture, correlation and marker localisation in high-resolution feedback^11^. As with every invention, the frame-based photosensor adopted in IFC can suffer from inherent constraints that hinder its performance in delivering high throughput results while maintaining rich spatial information.

Considering the limited field of view provided by microscopy or interrogation point in IFC, cells in the flow can only remain within the bounds of the scene for an extremely brief period. The frame-based capturing techniques applied can be vulnerable against consecutive fast-moving targets, especially in equipment with high frame intervals, high-velocity objects between two frames can lead to motion blur, ghost detection and other motion-induced artifacts^17^. Even though a professional-grade camera or sensor can be integrated to perform the image acquisition to reduce motion impact, the data redundancy caused by high frame-rate can obstruct the efficiency in data-intensive and delay-sensitive tasks^10^. Also, such a device’s expensive cost and maintenance can be overbearing for research and start-up projects. In terms of fluorescence analysis in IFC, which is an indispensable combination for revealing cell signalling, co-localisation, cell-to-cell interaction and DNA integrity in large-scale populations^11^. The dynamic range of current photosensors can have difficulties in perceiving limited fluorescence signals and overexcitation from light sources can lead to phototoxicity, photobleaching and tissue heating on cells^8^. These further exacerbate the trade-off relationship between speed, sensitivity and spatial resolution^6^. Increasing the throughput of IFC without jeopardising spatial resolution and sorting purity remains one of the outstanding obstacles in the field of IFC^7^. These inspired us to develop a data- and cost-efficient fluorescence-sensitive high-throughput neuromorphic cytometry to challenge these conundrums.

Neuromorphic vision is capable of detecting objects in motion by individually adapting brightness changes in each pixel and when there is no motion included in the field, pixels will remain inactivated^12^. This is in comparison to a frame-based pixel array, which is synchronously timed to a global shutter. The unique mechanism in neuromorphic allows real-time processing with low latency, leading to great application in object tracking, recognition and motion analysis^19^. As the maximum detection range of fluorescence signal is highly dependent on the dynamic range of the applied sensor^18^, the high dynamic range (*>* 120 dB) provided by an event-based photodetector can be an ideal candidate for visualising fluorescence-tagged objects, especially in low lighting or dark scenarios. To evaluate the possibility of integrating neuromorphic vision as a substitution for frame-based sensors in IFC and its feasibility of visualising microscale targets, the experiments were carried out with different sizes of polystyrene-based microparticles and THP-1 cells flowed within a microfluidic channel to create the essential dynamic contrast between targeted objects and the background for neuromorphic imaging. Herein, we are delivering an early demonstration of neuromorphic cytometry to perform object counting and size estimation on microscale particles and THP-1 cells, presenting the first instance of neuromorphic-enabled cell measurements and building upon our previous endeavour on introducing neuromorphic architecture as an alternative to overcome the conventional frame-related challenges in IFC.

### II. RELATED WORK

Imaging flow cytometry can offer rich spatial information in terms of cell interpretation; however, as with many imaging systems, IFC is also bound to the triangle of imaging constraints – speed, sensitivity and resolution. Increasing one of the parameters can lead to degradation in others^15^. In contrast, event-based cameras are more robust in handling low-lighting conditions and highly dynamic scenes owning to their asynchronous firing pixels, and the high-resolution event data can support over 3 *≥*s frame-rate^13^. Adopting an event-focused vision and architecture can significantly reduce the data redundancy and possibly elevate the performance in throughput and sensitivity without or with a minor reduction in resolution. He et al. ^4^ utilised the neuromorphic-enabled imaging classification and accomplished a mean average precision of 98.52 % at a speed of over 1000 fps and performed 3D reconstruction of the measured subjects via intensity and contour extraction. Moreover, the recent work conducted by Abreu et al. ^1^ utilising an event-based camera to conduct binary particle classification with a spiking neural network (SNN) and achieved 98.45 % testing accuracy, further consolidates the feasibility of implementing neuromorphic architecture in the field of cytometry. To the best of our knowledge, our previous work, Zhang et al. ^21^, is the first feasibility demonstration of conceptualising and building a *neuromorphic-enabled flow cytometry*, delivering early imaging of microscale objects under an event-based vision. All these endeavours indicated the potential of adopting a neurmorphic vision and architecture in delivering cell measurements, exhibiting a high level of accuracy and possible advanced throughput of neuromorphic cytometry.

## III. METHODOLOGY

### A. Sample preparation

Polystyrene-based microparticles were prepared in the initial testing to evaluate the performance of neuromorphic vision in capturing microscale targets. 10 wt.% concentrated 8 *μ*m and 15 *μ*m microparticle solutions were acquired (Sigma-Aldrich, USA) to emulate the relative size of human blood cells (erythrocyte, leukocyte) and abnormal cells (e.g., cancer cells). 8 *μ*m particle solution was diluted in 1:250 with deionised water to maintain a sufficient count and spacing between particles, preventing statistical inadequacy in sample size and possible aggregation. As the concentration of the particles was distributed by weight, 15*μ*m particle solution was diluted in 1:38 to obtain a similar number with diluted 8 *μ*m particles. Once the dilutions were completed, the sample particles were introduced into respective microfluidic channels to acquire neuromorphic feedback on counting and size estimation. To establish a baseline comparison, the same concentrations of the particles were prepared and analysed by a flow cytometer (CytoFLEX, Beckman Coulter) to obtain the measurement of counts and sizes to ensure the validity of the results.

Human monocytic cell line, THP-1 cells were maintained in a 37 °C and 5 % CO_2_ incubator and harvested in this experiment to examine the compatibility of neuromorphic vision with real sample cells and determine whether the complexity in cells can degrade or impact the visualisation and detection algorithm. The targeted cell line was prepared at the concentration of 1×10^6^ cells/mL. The cell solution was first centrifuged at a speed of 400 g for 5 minutes for sample separation. Then, the cell pellet was collected cells and re-suspended with a flow cytometry buffer made of 0.02 % sodium azide, 0.5 % bovine serum albumin (BSA) and 2 mM ethylenediaminetetraacetic acid (EDTA) in phosphate-buffered saline (PBS). The sample solution was vortexed and equally distributed into aliquots for neuromorphic analysis and CytoFLEX baseline comparison.

### B. Microfluidic imaging platform

A 60 *μ*m-height and 100 *μ*m-width microfluidic chip was fabricated utilising a standard photolithography protocol^20^. The chip contains one inlet and one output channel to create a basic delivery and visualisation pathway for the target samples. In addition, the channel employed a height of 60 *μ*m to focus the particles into a relatively limited plane to avoid a broad depth range and excessive calibration caused by it. In this platform, an event-based camera, Evaluation Kit 4 (EVK4, Prophesee, France) was integrated into a microscope (IX73 Inverted Microscope, Olympus, Japan) for proper magnification (10 ×) of the microscale objects and monitoring the dynamics of the field of view at the centre of delivering channel.

### C. Microparticle and cell experiments

The microparticles in the size of 8 and 15 *μ*m and THP-1 cells were adopted in this experiment to investigate the capacity of asynchronously activated pixels in tracking microscale objects. The diluted sample solution was loaded into a 1 mL syringe and distributed into the channel by actuation of a syringe pump (LEGATO® 200 Syringe Pumps, KD Scientific Inc, USA) at the flow rate of 10 *μ*L/min. As the objects travelled through the section of interest, the physical characteristics of the targets were collected and later analysed for object counting and size estimation by using the modified `metavision_psm.py` provided by Prophesee. For the baseline comparison, the same dilution factor was applied to the samples and analysed via CytoFLEX to provide outputs on total counts and estimated sizes.

## IV. RESULTS AND DICUSSION

The proposed schematic of our neuromorphic imaging flow cytometry is illustrated in Fig. 1 to demonstrate the working principle and expected future implementation. As the sample flow is introduced into the channel, the neuromorphic sensor will detect the ongoing subjects once it enters the view. The captured imaging will be fed to the processing centre for denoising, filtering, object counting, size and morphology estimation, determining whether the object is of interest and sending out a signal to the dielectrophoretic (DEP) sorter for sorting decisions. Once the DEF device receives the outcome, the electrodes will gently pull the targeted object into the pathway to the collection outlet. The unaltered objects will enter the waste outlet by default geometrical design.

**FIG. 1.**
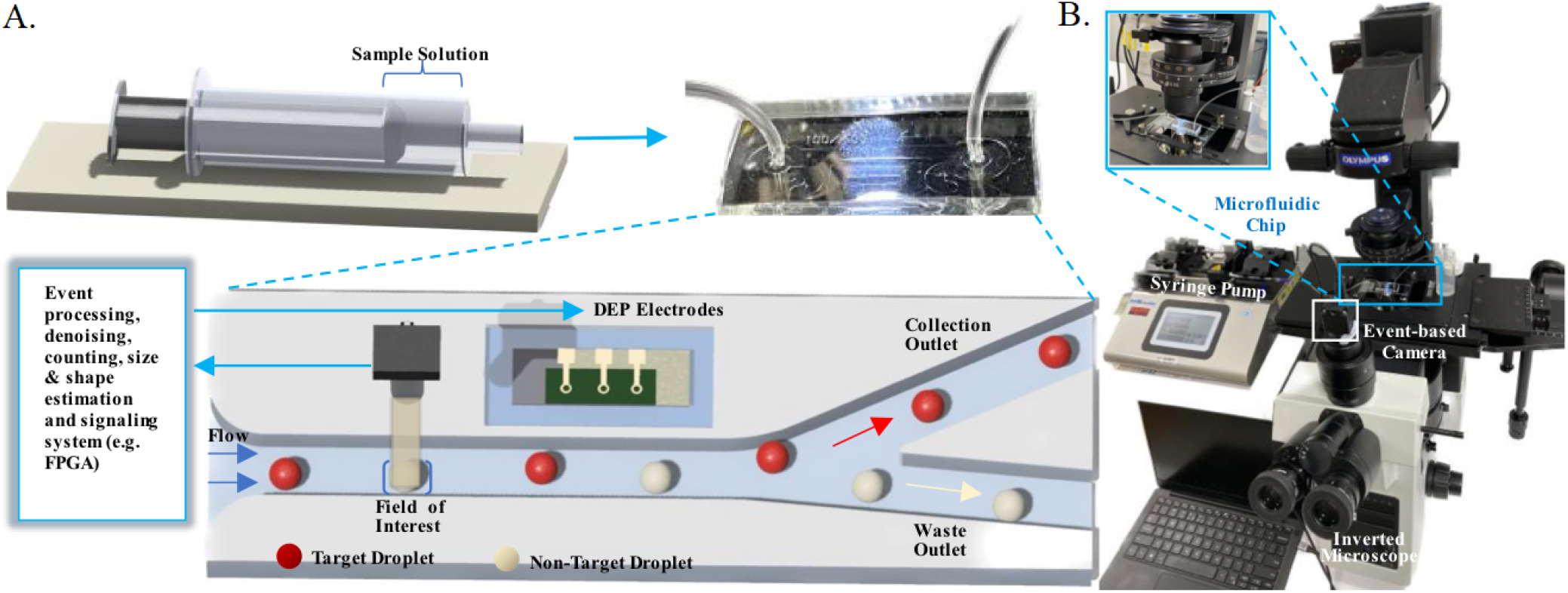
(A) Proposed schema of neuromorphic cytometry. The target samples were introduced into the microfluidic channel in a flow. Features of the samples were captured by an event-based camera, sending the inputs into the processing unit for classification and sorting purposes. After decision-making, the target object was directed into the collection channel via DEP sorter. (B) The current setup of the platform with an event-based camera integrated into a microscope for recording and capturing the sample flow.

One of the novel attempts that should be explored with neuromorphic sensors is the high dynamic range that is suitable for capturing fluorescence signals. Fig. 2. illustrates the documented throughput of fluorescence IFC with their respective dynamic range and the work published by Vinegoni et al. ^18^ highlighted that the maximum range of the fluorescence detected is majorly dependent on the dynamic range provided. Even though that recent efforts have achieved a high throughput with high-resolution imaging in detecting fluorescence-tagged objects, these works while remarkable, only adopted compensation strategies such as time-delay integration and virtual-freezing technique, the direct correlation and performance of high dynamic range in registering fluorescence signals remain unknown. To our knowledge, there hasn’t been an IFC that exploited such a high dynamic range (*>* 120 dB) compared to the neuromorphic camera, future endeavours on the subject can be extremely informative and promising in the aspect of developing next-generation IFC.

**FIG. 2.**
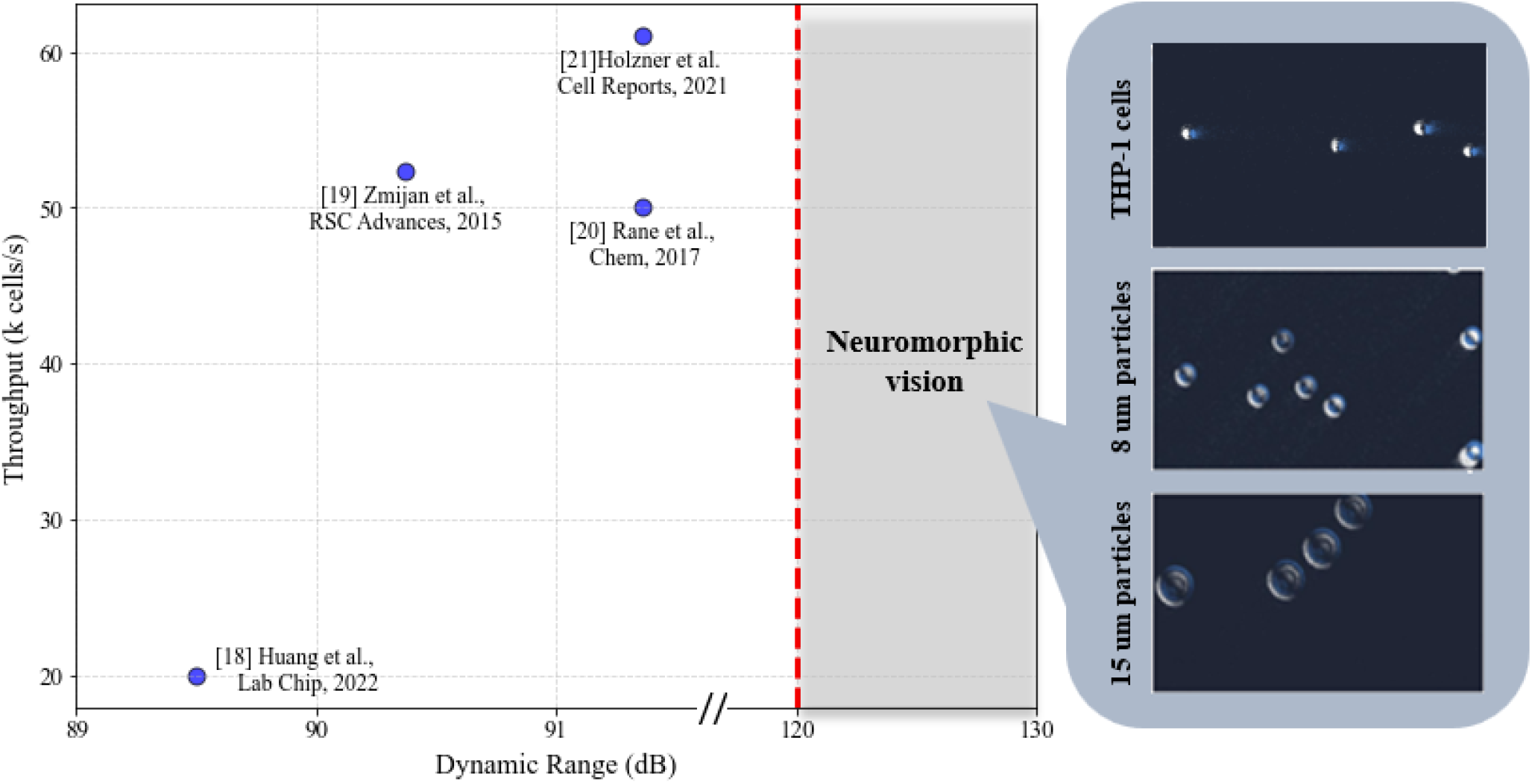
Performance of the previously reported fluorescence IFC with their incorporated dynamic range. The grey shade highlighted the unattained outputs and dynamic range provided by the neuromorphic vision and sample imaging of different microscale targets.

At the current stage, our platform can visualise and perform measurements on basic physical properties of microscale objects. In Fig. 3, the precise contour and relative size difference in neuromorphic view can be observed and compared to a phase contrast vision under a conventional microscope (ECLIPSE Ts2-FL, Nikon, Japan). Both imaging techniques were competent in delivering essential visual feedback without losing the key integrity of the subjects. In addition, when conducting microfluidic delivery with microscale targets, some of the travelling objects will inevitably adhere to the internal surface of the channel, rendering multiple imaging obstructions and deviating the focus from subjects. In neuromorphic, as the necessary movements are required to create the contrast between objects and the background, objects that remain in a static position will not trigger the activation of the pixels. This unique architecture can mitigate that consequential image effect caused by attached objects, significantly reducing the labour required for post-imaging processing and focusing on subjects of interest.

**FIG. 3.**
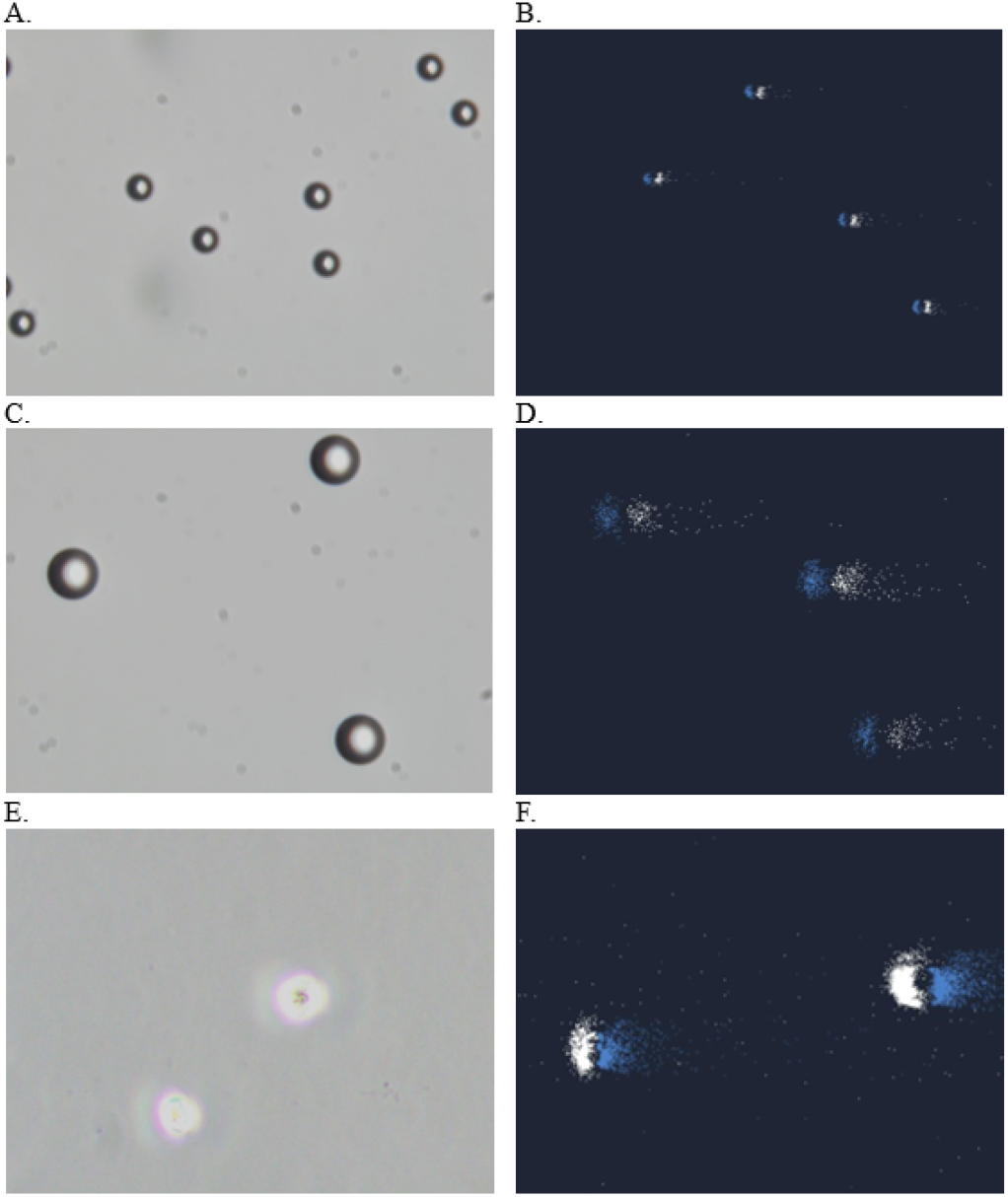
Conventional microscopic imaging and neuromorphic imaging of 8 *μ*m, 15 *μ*m microparticles and THP-1 cells. (A), (B) Microscopic and neuromorphic images of 8 *μ*m microparticles. The neuromorphic image on 8 *μ*m microparticles. (C), (D) Microscopic and neuromorphic images of 15 *μ*m microparticles. (E), (F) Microscopic and neuromorphic images of THP-1 cells.

With the implementation of object counting and size estimation, the number of microscale objects with their respective sizes can be estimated during the flow. Conventional cytometry scatter plots were provided in Fig. 4(A), (D), (G), including the gated population of 8, 15 *μ*m microparticles and THP-1 cells based on their sizes and granularity from FSC-A and SCC-A. For the 8 *μ*m microparticle solution, in Table I and Fig. 4(B), the total event count was 3189 with 95.6 % gated for 8 *μ*m microparticles, formulating a highly concentrated pattern of distribution around 1.1 M in FSC-A measurements with CytoFLEX; in Fig. 4(C), the neuromorphic detection was adopted and generated an overall count of 3086 with gating tolerance between ± 2 *μ*m in respective size, and yields 98.1 % of the population as 8 *μ*m microparticles. For the 15 *μ*m microparticle solution, in Fig. 4(E), the total event count was 3220 with 92.9 % gated for 15 *μ*m particles with CytoFLEX, presenting a concentrated distribution around 3.4 M in FSC-A measurement; in Fig. 4(F), the overall neuromorphic count was 3898 with 98.7 % gated as 15 *μ*m microparticles. Both technologies yield remarkable counting similarities and purities in terms of measuring pure polystyrene-based beads in different sizes, constructing an extremely alike size of distribution in describing the sample population. As the sample solutions are prepared with only microbeads, our strategy can compete with conventional cytometry in providing a higher purity and accuracy. For THP-1 cells, in Fig. 4(H), a total event number of 4429 was estimated by CytoFLEX with 40.6 % identified as live THP-1 cells, 26.6 % dead cells and 28.8 % debris; in Fig. 4(I), the neuromorphic count was 3056 and condition of the cells and division into subsets remain inaccessible owing to the inability to contrast different level of granularity. Although the majority of the ongoing objects in varied sizes can be detected, such discrepancy in the total count can be caused by the significant amount of tiny debris in the cell solution, which can be easily recognised as noise-like events during the flow in neuromorphic. As a result, our platform achieved high correspondence with commercialised cytometry when conducting measurements on pure microscale objects, however, in the context of cell measurements in real practice, the debris, doublets, complexity, viability and cycle of cells can raise the parameters to be implemented in order to distinguish various components in cell culture.

**TABLE I.** CytoFlex and neuromorphic output on the total event count of 8, 15 *μ*m microparticles and THP-1 cells in their sample solution.

| Subject name | Total population (CytoFlex) | Target population | Total population (Neuromorphic) | Target population |
| --- | --- | --- | --- | --- |
| 8 $\mu\text{m}$ microparticles | 3189 | 3049 (95.6 %) | 3086 | 3027 (98.1 %) |
| 15 $\mu\text{m}$ microparticles | 3220 | 2990 (92.9 %) | 3898 | 3848 (98.7 %) |
| THP-1 cells | 4429 | 1799 (40.6 %) | 3072 | N/A |

**FIG. 4.**
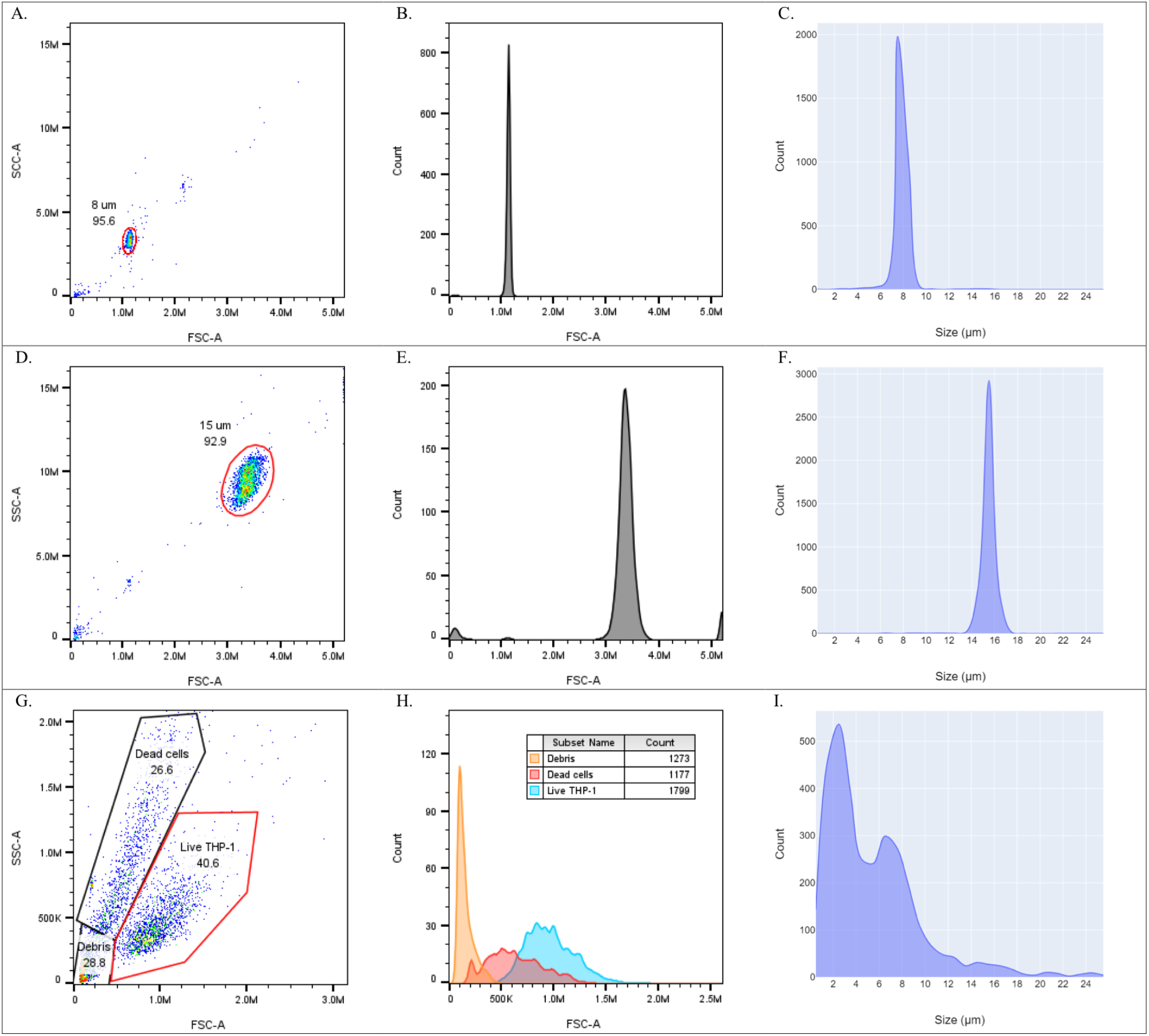
Cytometry and neuromorphic measurements on microparticles and THP-1 cells. (A) CytoFlex interpretation of 8 *μ*m microparticles based on forward scatter area (FSC-A) and sideward scatter area (SSC-A), gating 95.6 % of the population being 8 *μ*m microparticles. (B) Histogram of 8 *μ*m microparticle size distribution based on FSC-A measurement. (C) Histogram of 8 *μ*m microparticle size distribution based on neuromorphic outputs. (D) CytoFlex interpretation of 15 *μ*m microparticles based on FSC-A and SSC-A, gating 92.9 % of the population being 15 *μ*m microparticles. (E) Histogram of 15 *μ*m microparticle size distribution based on FSC-A. (F) Histogram of 15 *μ*m microparticle size distribution based on neuromorphic outputs. (G) CytoFlex interpretation of THP-1 cells based on FSC-A and SSC-A, gating 40.6 % as live THP-1 cells, 26.6 % dead cells and 28.8 % debris. (H) Size distribution of live THP-1 cells, dead cells and debris based on FSC-A. (I) Histogram of THP-1 cell size distribution based on neuromorphic outputs.

During the early stage of testing, MCF-7 cells were observed under the platform for testing the compatibility of neuromorphic vision with biological samples. In Fig. 5, in addition to the morphology acquisition, under phase contrast, the internal nucleus-like objects were identified with distinct clarity during the view without the aid of any staining techniques. Such a phenomenon can be supported by the high dynamic range provided by neuromorphic vision. However, as this is the first instance of the event, we cannot conclude an absolute explanation on the matter and further investigations and parallel studies on various cell lines are required to verify the specific mechanism behind it. Nevertheless, if the assumption is valid, this unique feature can contribute a deeper insight into neuromorphic cytometry and can be utilised as an additional metric in conducting cell measurement and sorting.

**FIG. 5.**
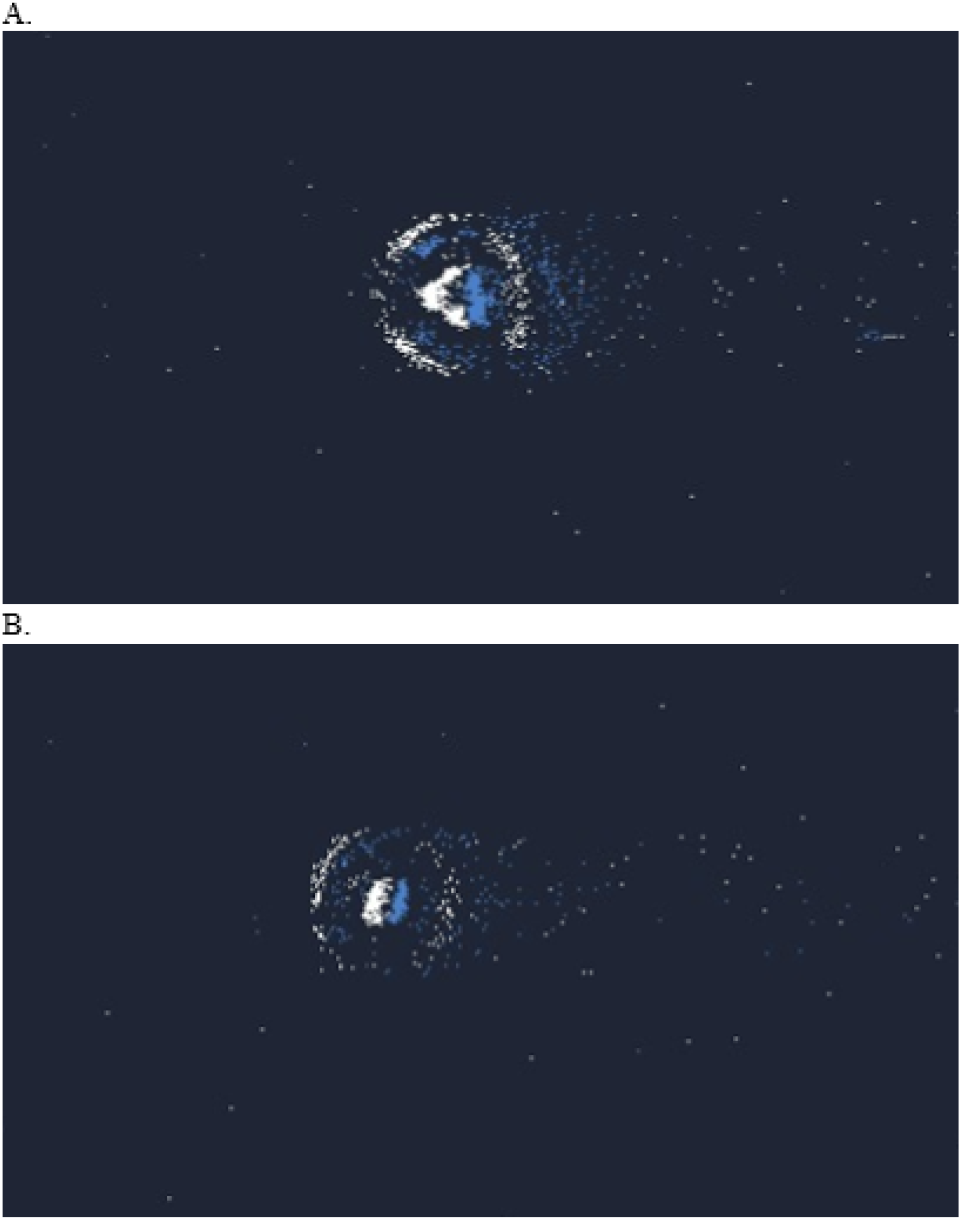
(A), (B) Nucleus-like objects in MCF-7 cells captured by neuromorphic vision under 20*×* magnification.

In this work, we have concluded our first milestone in performing cell population measurement in terms of object counting and size estimation. The high consistency and purity in measuring microbeads in different sizes compared to commercialised cytometry indicated the measurement accuracy and alike characterisation of neuromorphic cytometry. Although the function of classifying the cell condition and surrounding events are putting forward, the low latency and event-focused architecture in neuromorphic has the potential to outperformance the conventional IFC in throughput and data saving, enabling a prospect of a data- and cost-efficient fluorescence-sensitive high-throughput neuromorphic cytometry.

